# High-Throughput Image Analysis of Lipid-Droplet-Bound Mitochondria

**DOI:** 10.1101/2020.03.10.985929

**Authors:** Nathanael Miller, Dane Wolf, Nour Alsabeeh, Kiana Mahdaviani, Mayuko Segawa, Marc Liesa, Orian Shirihai

## Abstract

Changes to mitochondrial architecture are associated with various adaptive and pathogenic processes. However, quantification of changes to mitochondrial structures are limited by the yet unmet challenge of defining the borders of each individual mitochondrion within an image.. Here, we describe a novel method for segmenting Brown Adipose Tissue (BAT) images. We describe a granular approach to quantifying subcellular structures, particularly mitochondria in close proximity to lipid droplets, peri-droplet mitochondria. In addition, we lay out a novel machine-learning-based mitochondrial segmentation method that eliminates the bias of manual mitochondrial segmentation and improves object recognition compared to conventional thresholding analyses. By applying these methods, we discovered a significant difference between cytosolic and peridroplet BAT mitochondrial H_2_O_2_ production, and validated the machine learning algorithm in BAT via norepinephrine-induced mitochondrial fragmentation and comparing manual analyses to the automated analysis. This approach provides a higher-throughput analysis protocol to quantify ratiometric probes in subpopulations of mitochondria in adipocytes.

## INTRODUCTION

Image acquisition and analysis are critical tools for the quantification of mitochondrial parameters. Imaging data can provide not only morphological measurements (Joshi, Crouser, Julian, Schanbacher, & Bauer, 2000), but can also be used to quantify membrane potential (ΔΨ), mitochondrial mass, ROS production, and calcium concentrations, among others. A challenge facing the microscopist is the quantification of imaging data without biasing the results of the study. Automated image analysis provides an unbiased, repeatable, and high-throughput method to quantify high-resolution micrographs. The use of macros (a set of instructions for a computer to perform repeatedly) is the most amenable way to increase image analysis throughput with current tools (Mutterer & Rasband, 2012). A macro reduces the amount of time to analyze a single image while ensuring that subsequent images are analyzed in an identical manner.

Brown Adipose Tissue (BAT) provides a unique model for the study of mitochondria interaction with lipid droplets due to its high density of frequently overlapping mitochondria in the cytosol packed tightly around lipid droplets. Brown adipose tissue (BAT) is specialized to utilize energy to produce heat upon activation by norepinephrine (Wikstrom et al., 2014; Zingaretti et al., 2009). This specialization depends on changes to mitochondrial dynamics. Cell biologists have hypothesized (Wikstrom et al., 2007) that mitochondria may specialize into different populations within a cell, but evidence for this hypothesis is rare due to the difficulty of segmentation for analysis. Previous publications has supported this hypothesis in muscle, (Glancy et al., 2015) but other cell types remain to be analyzed in such a way. For instance, a question of interest is: How different are peridroplet mitochondria from their cytosolic counterparts? Recent work has shown that peri-droplet mitochondria are distinct from cytosolic mitochondria (Benador et al., 2018). Previous work in other cell types has focused on quantifying the entire network or manual classification of network morphology (Chaudhry, Shi, & Luciani, 2019; Cribbs & Strack, 2009; Leonard et al., 2015; Mahdaviani et al., 2017; Valente, Maddalena, Robb, Moradi, & Stuart, 2017). In contrast, our study focuses on direct quantification of individual mitochondria. Improvements in computer processing power and graphical handling enable a novel solution to this difficult problem: machine learning. Machine learning is the process by which humans train a computer program to perform a particular task. The machine learning algorithm utilized a random forest model and is one of many iterative machine learning algorithms (Breiman, 2001).

Due to considerations of cost and transparency, the open-source programs ImageJ (https://imagej.nih.gov/ij/) and its plugin-rich counterpart FIJI (Schindelin et al., 2012) (http://fiji.sc/) are utilized for these analysis protocols and data. FIJI provides several plugins not available in stock ImageJ that greatly expedite analysis. Chief among these tools is the WEKA trainable segmentation plugin (Arganda-Carreras et al., 2017), which enables rapid training and deployment of machine-learning segmentation protocols. Tools of this nature are incredibly resource intensive and, as such, must be used on computers with sufficient processing and graphical power. Both ImageJ and FIJI are open-source and widely documented, providing ideal tools for scientific use.

Similar tools have previously been utilized to quantify mitochondria, albeit in a more limited manner. Koopman *et al*. introduced the concept of using a machine learning classifier to morphologically characterize and bin mitochondria (Koopman, Visch, Smeitink, & Willems, 2006). This technique has been further refined by other groups (Leonard et al., 2015; Valente et al., 2017). Both of these approaches did not fully quantify mitochondrial morphology, but rather quantified the population of each type of mitochondrion, e.g. tubular vs. punctate. Previous work has also described what parameters best define mitochondrial morphology (Joshi et al., 2000). While the conventional approach of manual thresholding for segmentation is valid, it is subject to the analysts’ bias (Cribbs & Strack, 2009). Combining these above approaches – using machine-learning to segment mitochondria which are then individually quantified using established parameters – provides the most robust and accurate quantification to date. Software exists to perform a similar segmentation routine (Harwig et al., 2018), but it is not easily adaptable as an ImageJ script, while the methods provided here are cross-platform and usable within ImageJ or FIJI. Additionally, the data outputs from these two methods are not identical and may in fact complement each other.

## MATERIALS AND METHODS

### Primary brown adipocytes culture

Primary brown adipocytes (BA) were generated by differentiating pre-adipocytes isolated from BAT as described in detail previously(Assali et al., 2018; Benador et al., 2018; Cannon & Nedergaard, 2001; Wikstrom et al., 2014). BAT was harvested from 3 to 4-weeks-old WT and NCLX KO mice. In brief, The tissue was dissected from interscapular, subscapular, and cervical regions, minced, and transferred to a collagenase digestion buffer (2 mg/mL Collagenase Type II in 100 mM HEPES, 120 mM NaCl, 4.8 mM KCl, 1 mM CaCl_2_, 4.5 mM Glucose, 1.5% BSA, pH 7.4) at 37°C under constant agitation for 30 min. Collagenase digestion was performed in 37°C water incubator under constant agitation for 25 min with vortex agitation every 5 min. Digested tissue was homogenized and strained through 100 mm and 40 mm strainers. Cold DMEM was added to tissue digest and centrifuged twice (the last included washing and resuspension in new DMEM at 200 × g speed for 12 min at 4°C). Finally, cell pellets (preadipocytes) were re-suspended 5 mL growth medium (DMEM supplemented with 20% newborn calf serum (NCS), 4 mM Glutamine, 10 mM HEPES, 0.1 mg/mL sodium ascorbate, 50 U/mL penicillin, 50 mg/mL streptomycin) and plated in 6-well plates (Corning). Cells were incubated in 37°C 8% CO_2_ incubator. 24 h after isolation, the cells were washed to remove debris and medium was replaced. 72 h after isolation the cells were lifted using STEMPro Accutase, counted, and re-plated in differentiation media (growth media supplemented with 1 mM rosiglitazone maleate and 4 nM human recombinant insulin). Cells were differentiated for 7 days and medium was changed every other day. For transduction experiments, cells were transduced with the roGFP virus in differentiation day 0-3.

### Viral Transduction of ROS reporters

Mitochondrially-targeted Orp-1 roGFP (Morgan, Sobotta, & Dick, 2011) was adenovirally transduced into primary BAT isolated from 12-16 week old C57BL/6J mice overnight at a MOI of 2000 particles per cell. This resulted in >90% transduction. After an overnight incubation at 37°C and 8% CO_2_, the cells were given fresh media without viral particles and cultured for 72 hours prior to imaging.

### Microscopy

Images of BAT were obtained on a Zeiss LSM 710 or LSM880 microscope equipped with a plan-apochromat 100X (NA=1.4) oil immersion objective. Images were taken with a digital zoom of 1 or above. Images were at least 1024 × 1024 px.

## RESULTS

### Validation of machine learning classifier

The WEKA segmentation classifier was validated for BAT against human recognition of objects. Briefly, a human created ROIs manually around each mitochondrion in a cell, with 50-200 mitochondria per cell. These individual regions were quantified identically to the WEKA-generated or threshold-generated ROIs of the same cells. All three analyses were performed by the same individual, who manually thresholded at a level that facilitated object recognition, typically 10% of the maximum fluorescence intensity. All images were high resolution images of single cells cropped from a larger field of view. This cell-level analysis facilitates granular analysis of the data and examination thereof on a cell by cell basis.

### Training images used for WEKA training set

Images used for training of the classifier were high resolution and super resolution micrographs. Training was performed on manually cropped individual cells from different fields of view, more than 4 independent experiments, and on 2 different fluorescence markers of mitochondria: mitotracker green and Tetramethylrhodamine, Ethyl Ester, Perchlorate TMRE. This broad training set increased accuracy and flexibility of segmentation. Input images for segmentation should be of as high resolution as possible, but maximum pixel size utilized for analysis was 0.07 μm per px. Some enhancements can improve poor quality images for analysis, such as a previously published filtering method utilizing a median filter (Smith, Kovats, Lee, & Cano, 2006) or utilizing built in background subtraction in ImageJ, such as the rolling ball algorithm. Regardless of quality of input images, Figures 3 and 4 illustrate lower-quality, lower-resolution (0.165 μm per px) can still be used for this analysis.

### Mitochondria morphological parameters that describe mitochondrial networking

Multiple sources (Joshi et al., 2000; Koopman et al., 2006; Leonard et al., 2015; Molina et al., 2009; Nguyen, Beyersdorf, Riethoven, & Pannier, 2016; Twig et al., 2006; Wikstrom et al., 2007) have demonstrated that mitochondrial morphology can be described by various shape descriptors (Table 1). Chief among these shape descriptors is the circularity measurement (Circ), which compares the perimeter of the region of interest to the perimeter of a perfect circle of the same area. This measurement is primarily used to describe branching of the network, because it equals 1 when measuring a perfect circle, and < 1 for a starfish shape. Form factor (FF) is another frequently used shape descriptor, and is simply the inverse of circularity, meaning a starfish shape has a FF much greater than 1, while a perfect circle still has a FF of 1. Another frequent shape descriptor used for analysis of mitochondrial morphology is the aspect ratio (AR). AR measures the ratio of the long axis length of the ROI to the short axis length, and is a measure of elongation of the object. Solidity measures the concavity of the ROI, which can also be interpreted as branching or connectivity. Utilizing these parameters, mitochondria may be described in morphological as well as fluorescent detail.

**Table 1.** Common mitochondrial morphology parameters.

| Parameter | Information | Units | Notes |
| --- | --- | --- | --- |
| Fluorescence Intensity (FI) | Brightness of fluorophore | A.U. | Can be sum or average of all intensity for entire region. Ratios of multiple fluorophores are possible |
| Area (A) | Cross-sectional area of region | Pixels (px), $\mu\text{m}^2$ | |
| Perimeter (P) | Distance around region | Pixels (px), $\mu\text{m}$ | |
| Circularity (Circ) | Branching<br>$\frac{4\pi(\text{Area})}{(\text{Perimeter})^2}$ | A.U. | $0 < \text{Circ} \leq 1$;<br>Circ is 1 for a perfect circle; less than 1 for branched structures<br>$\text{Circ} = \text{FF}^{-1}$ |
| Form Factor (FF) | Branching<br>$\frac{(\text{Perimeter})^2}{4\pi(\text{Area})}$ | A.U. | $1 \leq \text{FF} < \infty$;<br>FF is 1 for a perfect circle; more than 1 for branched structures<br>$\text{FF} = \text{Circ}^{-1}$ |
| Aspect Ratio (AR) | Elongation<br>$\frac{\text{Long Axis}}{\text{Short Axis}}$ | A.U. | AR measures elongation, not branching |
| Solidity | Concavity<br>$\frac{\text{Object Area}}{\text{Convex Hull Area}}$ | A.U. | Creates a fully convex shape (hull) surrounding the object and measures the ratio of the object area to the convex hull area |

### Segmentation of cytosolic mitochondria versus peridroplet mitochondria reveals significant differences in H_2_O_2_ production

The analysis of these images proved to be challenging – how does one segment mitochondria in the cytosol apart from mitochondria immediately surrounding lipid droplets? By selectively labeling lipid, we can visualize the droplets in order to segment the mitochondria around them. While this approach is somewhat limited in its morphological analysis, an additional level of sub-segmentation could be added in order to separate each mitochondrion from its neighbors and measure each one’s morphology in addition to measuring the average fluorescence intensity of the whole population. Here, we demonstrate a novel segmentation method for BAT mitochondria using a lipid label for the analysis.

The original image contains 4 channels: Nile red (lipid), ORP1-roGFP (redox status, 2 channels), and bright field (BF). The BF channel is immediately discarded upon beginning analysis; it is a quality control channel included for manual review. The roGFP probe is a redox sensitive GFP fused to ORP1, which is itself a H_2_O_2_-specific antioxidant enzyme from yeast, making this probe a tool to measure H_2_O_2_ production in the mitochondrial matrix (Morgan et al., 2011). Nile Red is a lipophilic dye that marks lipid compartments selectively; it is used to locate and segment lipid droplets within BAT cells.

In order to evaluate cytosolic versus peridroplet mitochondria, the first step is to segment the lipid regions (Fig 1), which is accomplished with a basic automated thresholding step (Otsu method). This binarized image is then used for several other operations within the analysis. First, it is used to subtract the small amount of fluorescence bleed-through from Nile red into the reduced roGFP channel. Second, it is used as the seed for peridroplet-mitochondria recognition. This segmentation is accomplished by dilation of the lipid region *n* times, where *n* is user-defined at the outset of analysis. This step is user-defined because of several considerations, primarily due to artifacts resulting from diverse input images. Because images may be taken at different magnifications and metadata is not always present to convert pixel units to SI units, allowing the user to define the peridroplet region in pixel units removes potential bias and bugs from the analysis itself while providing extensive customization for various inputs. After dilation and automated segmentation, the region immediately surrounding the lipid droplet is measured.

**Figure 1.**
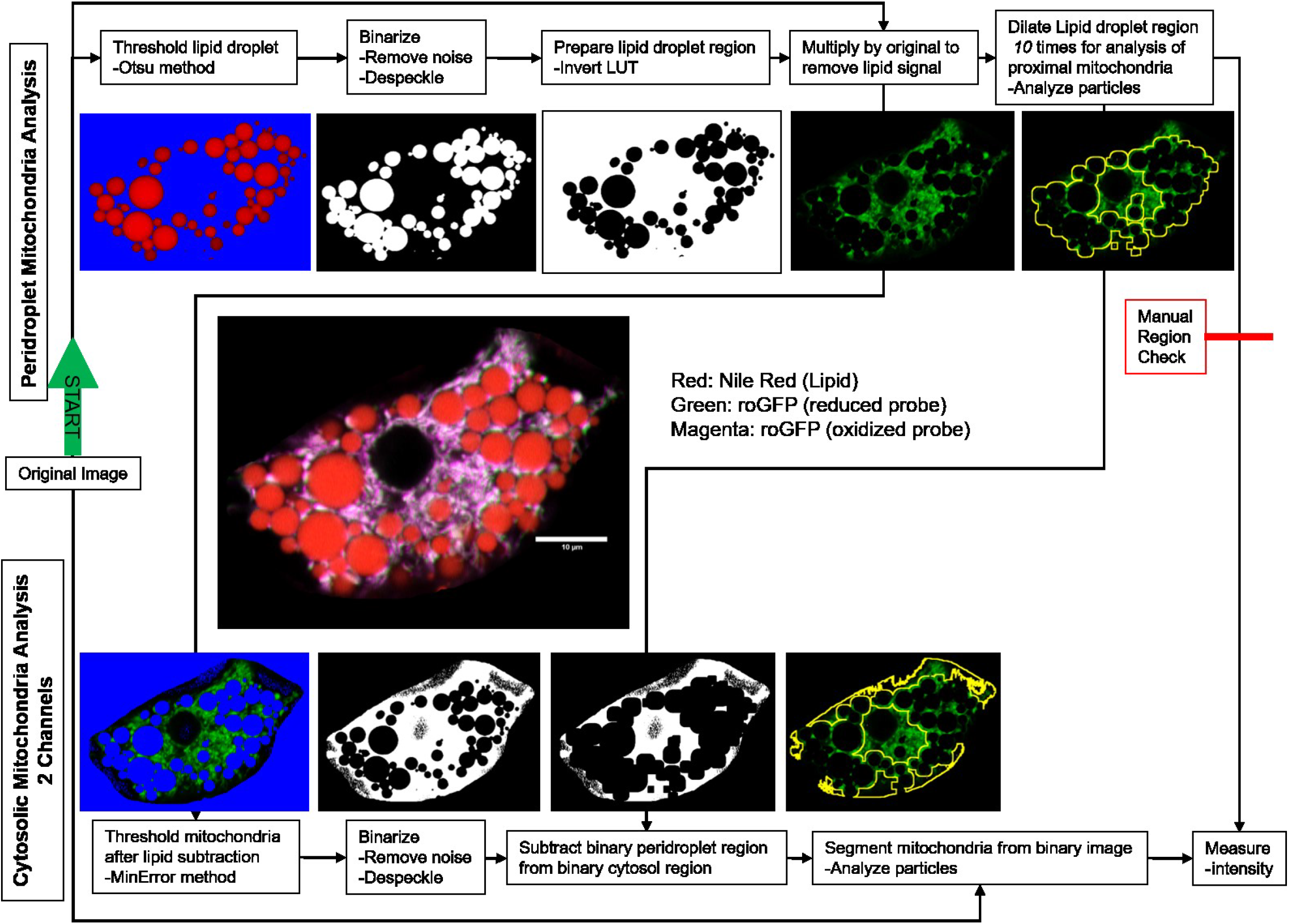
Schematic representation of quantification of peridroplet and cytosolic mitochondria in Brown Adipose Tissue. The original mutli-channel image is split into its component channels, which are then filtered for analysis. The lipid droplet is labeled with Nile Red dye, and this image region is used for recognition and segmentation throughout the analysis. The Nile red image is binarized and used multiple times in the analysis. First, it is used to subtract fluorescence bleed-through in the roGFP channel from the Nile Red. Then, the lipid droplet region is dilated *n* times, with *n* being empirically determined by the user to encompass peridroplet mitochondria. This dilated lipid region is then used to quantify the mitochondrial signal contained within. The cytosolic analysis builds on the previous steps, utilizing the dilated lipid region to remove any mitochondrial fluorescence in that region from the cytosolic image. The cytosolic image is then binarized, segmented, and measured similarly to the peridroplet image. Scale bar shown in original image is 10μm and is intentionally left out during analysis. Manual steps are outlined in red.

After the peridroplet region is measured, the dilated lipid image is used to subtract all peridroplet fluorescence from the cytosolic image, leaving only objects not analyzed in the previous step. This cytosolic image is then automatically thresholded using the MinError method in FIJI, then segmented using the “analyze particles” tool. This segmented image is then measured for the cytosolic data. Once these measurements are saved as a .csv file, they can be handled like other data arrays in various programs.

When BAT images were segmented to separate subcellular populations of mitochondria, we found that cytosolic mitochondria produce significantly more H_2_O_2_ than peridroplet mitochondria (Fig 2). The addition of menadione, a mitochondrial toxicant that induces ROS formation via redox cycling (Criddle et al., 2006) was used as a positive control and elicited a significant increase in H_2_O_2_ production in treated cells.

**Figure 2.**
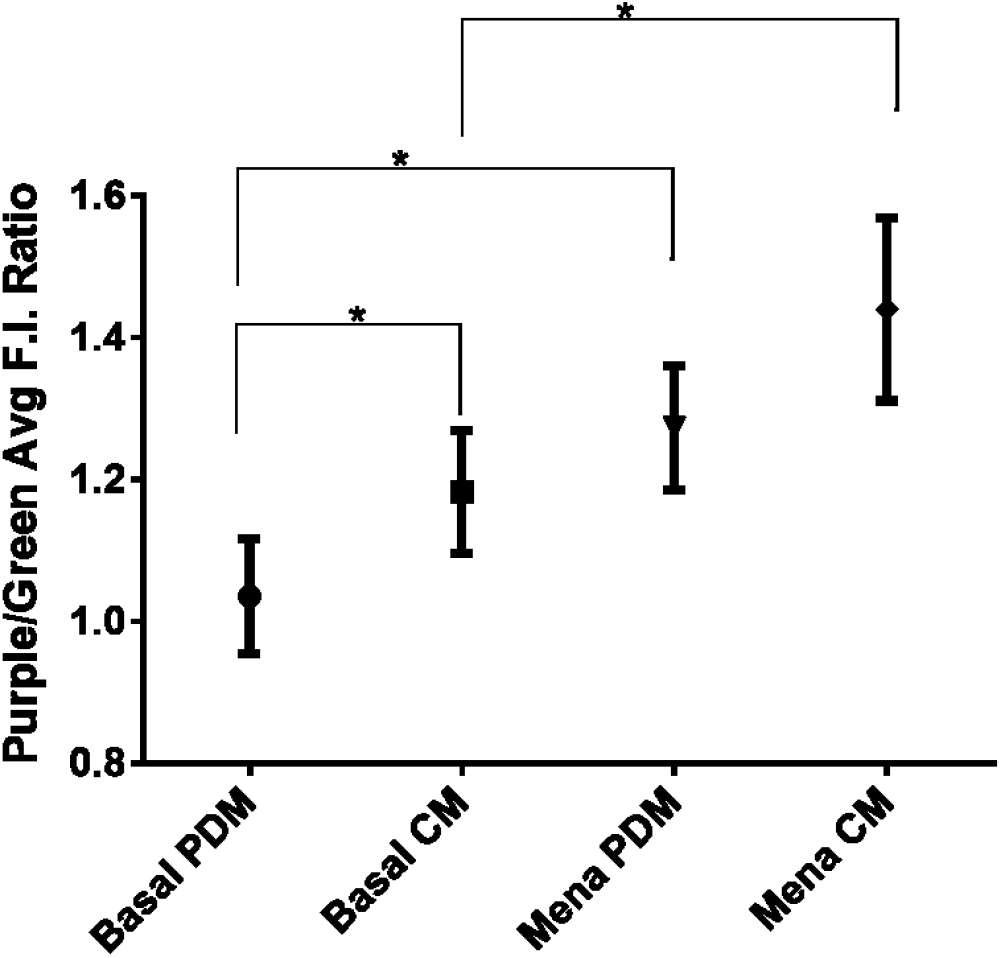
BAT mitochondria produce different amounts of H_2_O_2_ depending on their localization in relation to lipid droplets. BAT cytosolic mitochondria (CM) produce significantly more H_2_O_2_ than their peridroplet (PDM) counterparts. Addition of menadione increased ROS production significantly in both mitochondrial populations.

### Trainable WEKA segmentation is significantly more effective at mitochondrial segmentation compared to conventional thresholding methods

Trainable WEKA segmentation is a plugin for FIJI that enables machine-learning-based image segmentation. Previously, work has focused on classifying and counting mitochondria using machine learning (Leonard et al., 2015). Frequently, these approaches only provide counts of objects contained in each class. With these known limitations, we set out to 1) improve mitochondrial segmentation for analysis and 2) utilize machine learning in a novel way to that end.

Machine learning is a complex field with multiple algorithmic approaches, each with tradeoffs between speed, accuracy, processor utilization, and other system requirements. The FastRandomForest (Breiman, 2001) algorithm attempts to mitigate some of the system requirements and speed up computation, which is part of why it is commonly used in this context. To the user, the algorithm is mostly a backend affair, and once selected is not seen again during training. WEKA provides a graphical user interface (GUI) useful for training a classifier. Within this GUI, the user defines the number and names of each class to be trained (in this example, the two classes are “mitochondria” and “background”). After defining and naming classes, the user trains the classifier, which entails using any of FIJI’s selection tools (line, rectangle, circle, etc.) to define regions of an image belonging to a certain class. In Figure 3, this iterative process is demonstrated while training the classifier to distinguish mitochondrial signal from background noise. Figure 3F represents the final classification of the image, showing mitochondria in red and background in green. The final set of instructions for segmentation are stored in a file called a classifier with extension “.model” This published classifier is the product of over 100 training images containing tens to hundreds of discrete data points and used for subsequent analysis of new image sets (Fig 4).

**Figure 3.**
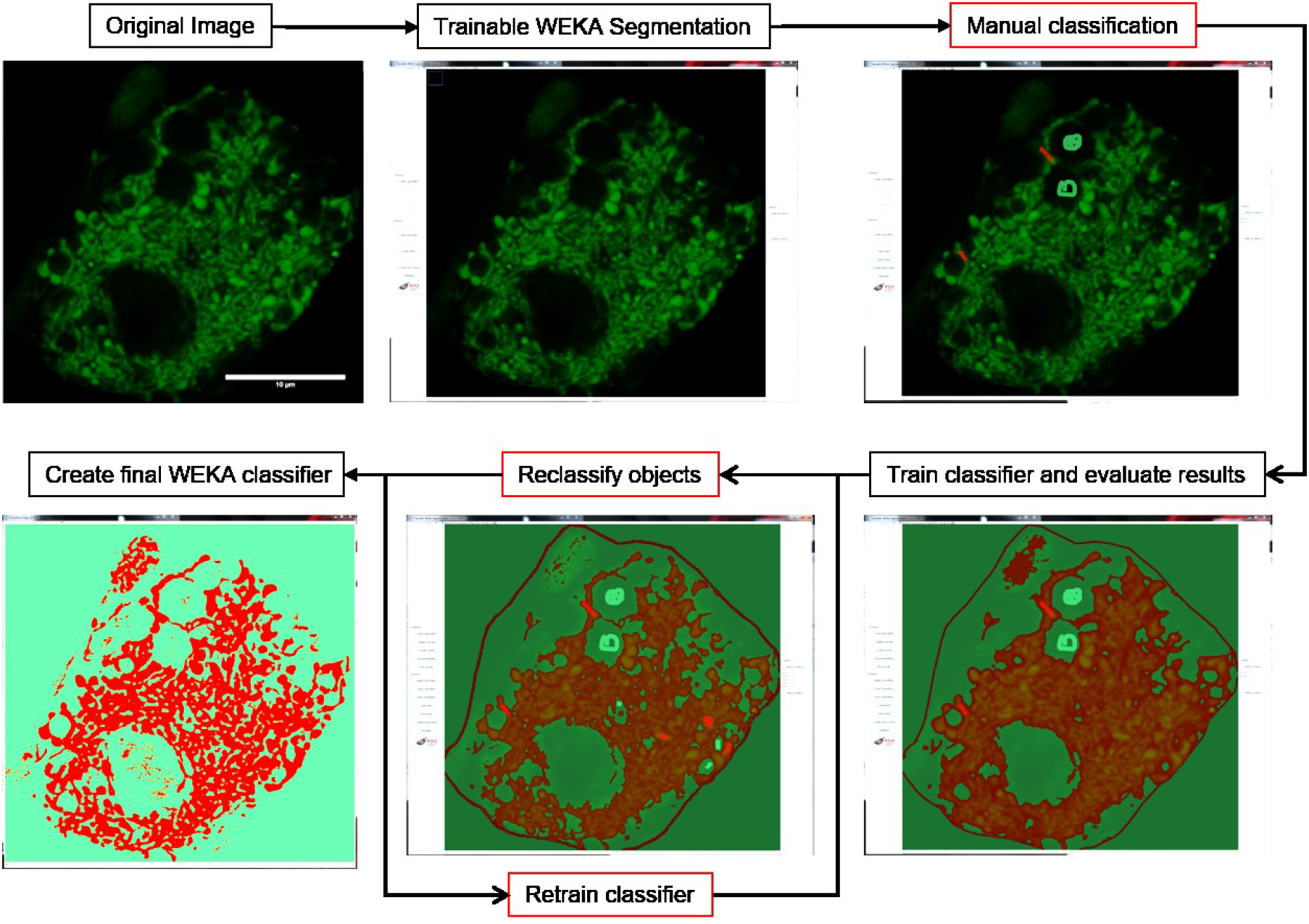
Representation of iterative training of the machine learning classifier using the WEKA trainable segmentation plugin for FIJI. Once an image is opened in FIJI, WEKA trainable segmentation can be called from the plugins menu. WEKA is capable of using several machine learning algorithms, with the most fitting being FastRandomForest considering time and processor utilization. Options set within the WEKA GUI can control the parameters used for machine learning, though several of these require additional plugins. Once open, the user defines the number and name of each class of object. Once the classes are defined, the user can use any selection tool available in FIJI (line, rectangle, circle, etc.) to define what image objects belong to which class. Upon clicking “train classifier,” the computer then calculates the best segmentation algorithm and returns a map of the classes overlaid over the training image which can be used to refine the class segmentation with subsequent rounds of assignment and training. Once WEKA returns a satisfactory segmentation of the image, the user can finalize the classifier to be used on new data sets. Scale bar shown in original image is 10μm and is intentionally left out during analysis. Manual steps are outlined in red.

**Figure 4.**
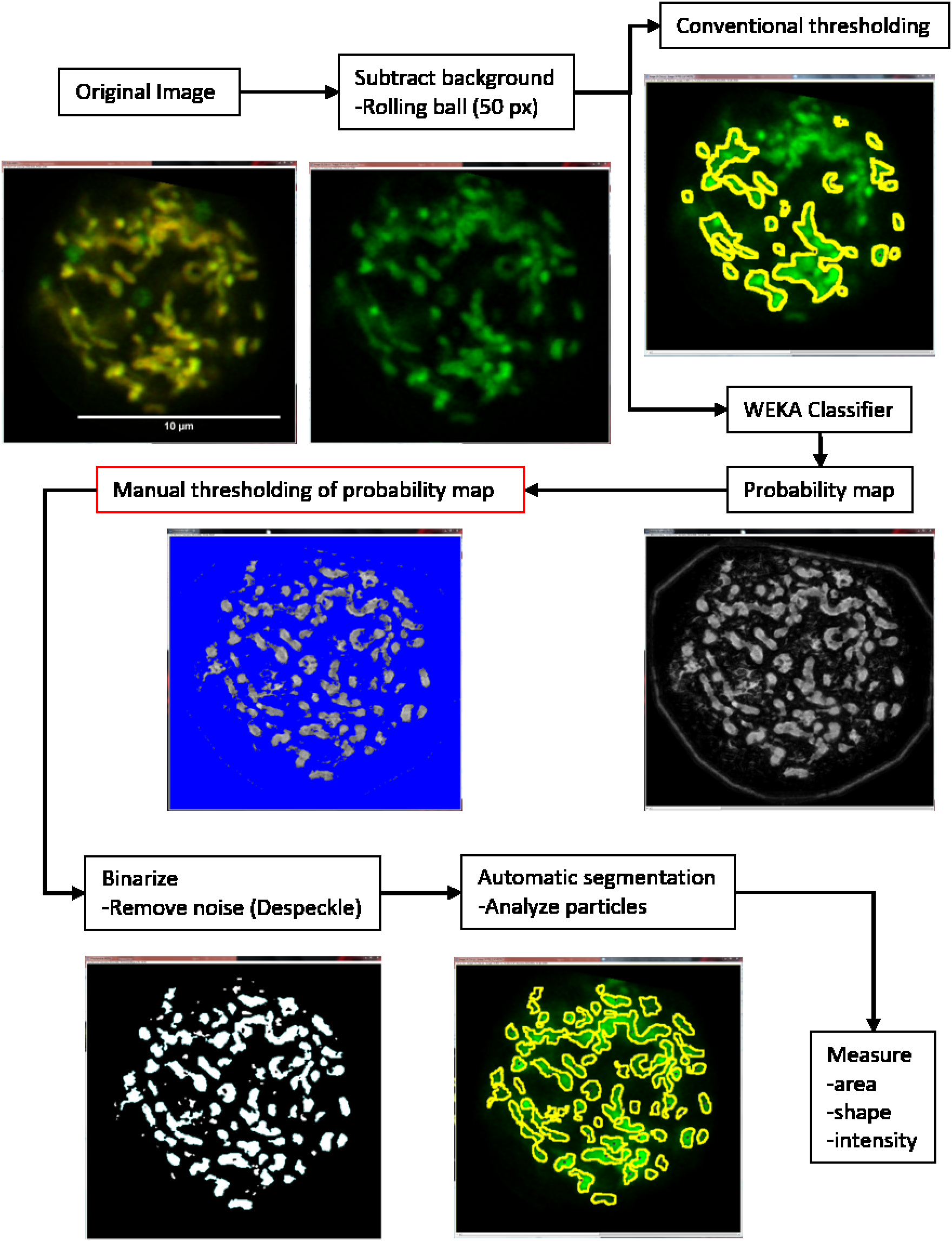
Workflow for a WEKA-based mitochondrial segmentation and analysis. The original image is initially filtered to remove uneven background using the rolling ball algorithm with a diameter of 50 pixels. This facilitates better recognition of mitochondria by the WEKA classifier. Shown at this branch point for comparison is traditional threshold-based segmentation of mitochondria in the same image. Once the image is classified with the WEKA classifier, it returns a probability map. A probability map is the algorithm’s best guess at which class each pixel in an image belongs to, in this case mitochondria or background, with a probability from 0 to 1. The user manually thresholds the certainty of the classification for best mitochondrial segmentation (though this step may be automated with validation). After the thresholding step, the image is binarized, the mitochondrial objects segmented for measurement, and then measured. Scale bar shown in original image is 10μm and is intentionally left out during analysis. Manual steps are outlined in red.

Because manual analysis of hundreds of images is time consuming, the WEKA classifier needed to be incorporated into a high-throughput analysis method. The analysis workflow into which it was incorporated is shown in Figure 4. WEKA segmentation provided segmentation of mitochondria with closer ROIs to the real objects (with less effect of background signal), as well as better separation of almost-touching objects compared to conventional filtering and thresholding methods. The output of the classifier is a probability map: an image where the pixel intensity value represents the computer’s certainty from 0 to 1 that a specific pixel belongs to a certain class (in Figure 4, the “mitochondria” class is shown). This probability map can be manually or automatically thresholded for certainty level, but adding a manual step at this point ensures data quality with a human checkpoint evaluating the classifier output.

When INS-1 cells were segmented with WEKA segmentation, the number of detected mitochondria increases twofold (Figure 5A). This increase is attributable to the more sensitive and specific detection of background between proximal mitochondria compared to conventional thresholding, resulting in fewer mitochondria being combined when in fact they are discreet objects. Because of the increased separation of mitochondria, mitochondrial perimeter is significantly decreased in WEKA-segmented measurements (Figure 5B). This decreased perimeter is the result of WEKA segmenting mitochondria while ignoring much of the airy fluorescent haze surrounding the mitochondria. Due to this improved segmentation, aspect ratio is increased (Figure 5C), circularity is decreased (Figure 5D), and solidity is decreased (Figure 5E). As an analogy, if outlining an object left a 1mm gap between the object and its outline, the object would be measured as being more round (circularity approaching 1, aspect ratio approaching 1, and solidity approaching 1). However, if the outline is more accurate and leaves only a 0.1mm gap between the object and the outline, the outline’s shape will more closely match the shape of the object, and its measurements would diverge from 1 because the object would be measured as less round. This increased accuracy of segmentation is illustrated in Figure 4 by comparing the traditionally thresholded image to the WEKA-segmented image.

**Figure 5.**
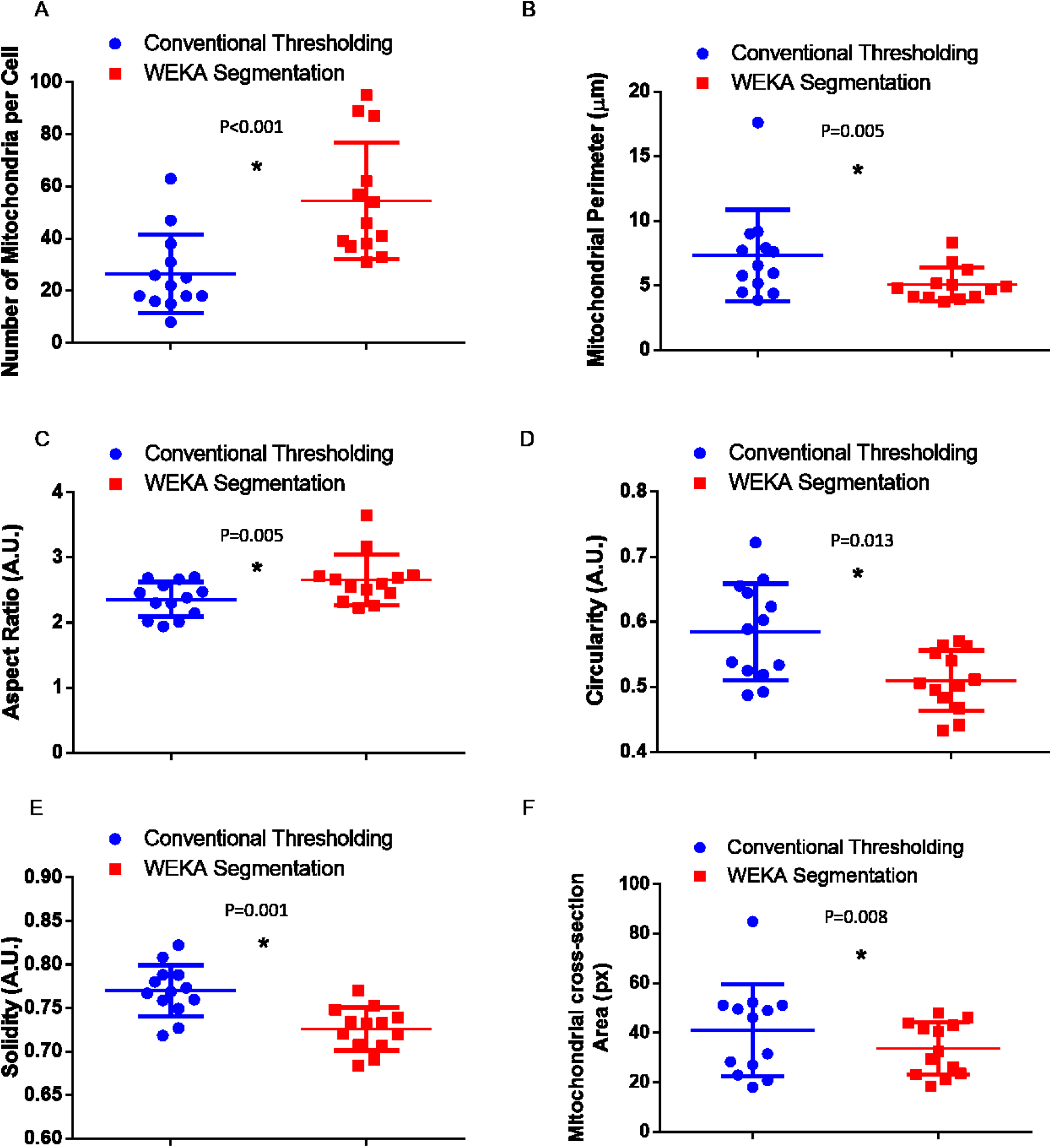
Comparison of mitochondrial morphological measurements segmented with WEKA segmentation versus traditional thresholding. WEKA segmentation is significantly better at recognizing, separating, and quantifying individual mitochondria (A). Because of this more accurate segmentation, the mitochondrion is measured much more closely to its actual boundaries instead of the airy haze surrounding it. As a result, the Perimeter readout is significantly reduced (B). This more accurate segmentation provides more accurate measurements of Aspect Ratio (C), Circularity (D), and Solidity (E), and cross-sectional area (F). Better object recognition leads to improved dynamic range of measurements. Each point represents an individual image, error bars SEM. * p≤0.05 by t-test.

### Detection of NE-induced mitochondrial fragmentation in BAT validated WEKA segmentation

BAT mitochondria fragment upon stimulation with NE (Wikstrom et al., 2014). We used this model to validate WEKA segmentation. WEKA segmentation was compared to the thresholding approach and to manual segmentation of mitochondria by ROI drawing. While manual analysis exhibited the largest dynamic range (1.3 AR, 0.31 Circ for Manual vs 0.008 AR, 0.08 Circ for thresholding) and provided the most robust results (Fig 6), which is to be expected, thresholding provided the least robust results – it is hampered by the background-increasing airy haze surrounding objects inherent to confocal microscopy. WEKA segmentation yielded a large decreases in mitochondrial connectivity (Circ and solidity), while also demonstrating a decreasing trend in AR upon fragmentation, with an AR of 2.0 (elongated) reducing to 1.7 (fragmented). This trend in AR is mitigated by an artifact inherent to the WEKA classifier included herein. Due to the inherent difficulty in mitochondrial segmentation, the classifier splits larger objects into smaller ones, such as the very elongated filamentous mitochondria in non-stimulated BAT. However, this bias toward splitting is necessary to retain mitochondrial recognition, as evidenced by WEKA segmentation’s trend to detect more mitochondrial objects. While it may be possible to reduce the impact of this over-segmentation artifact by utilizing a BAT image training set rather than a mixed image training set, it would also decrease the flexibility of the classifier. Alternatively, due to the higher throughput of analysis enabled by the WEKA classifier and macro, larger sample sizes would also offset this artifact.

**Fig 6.**
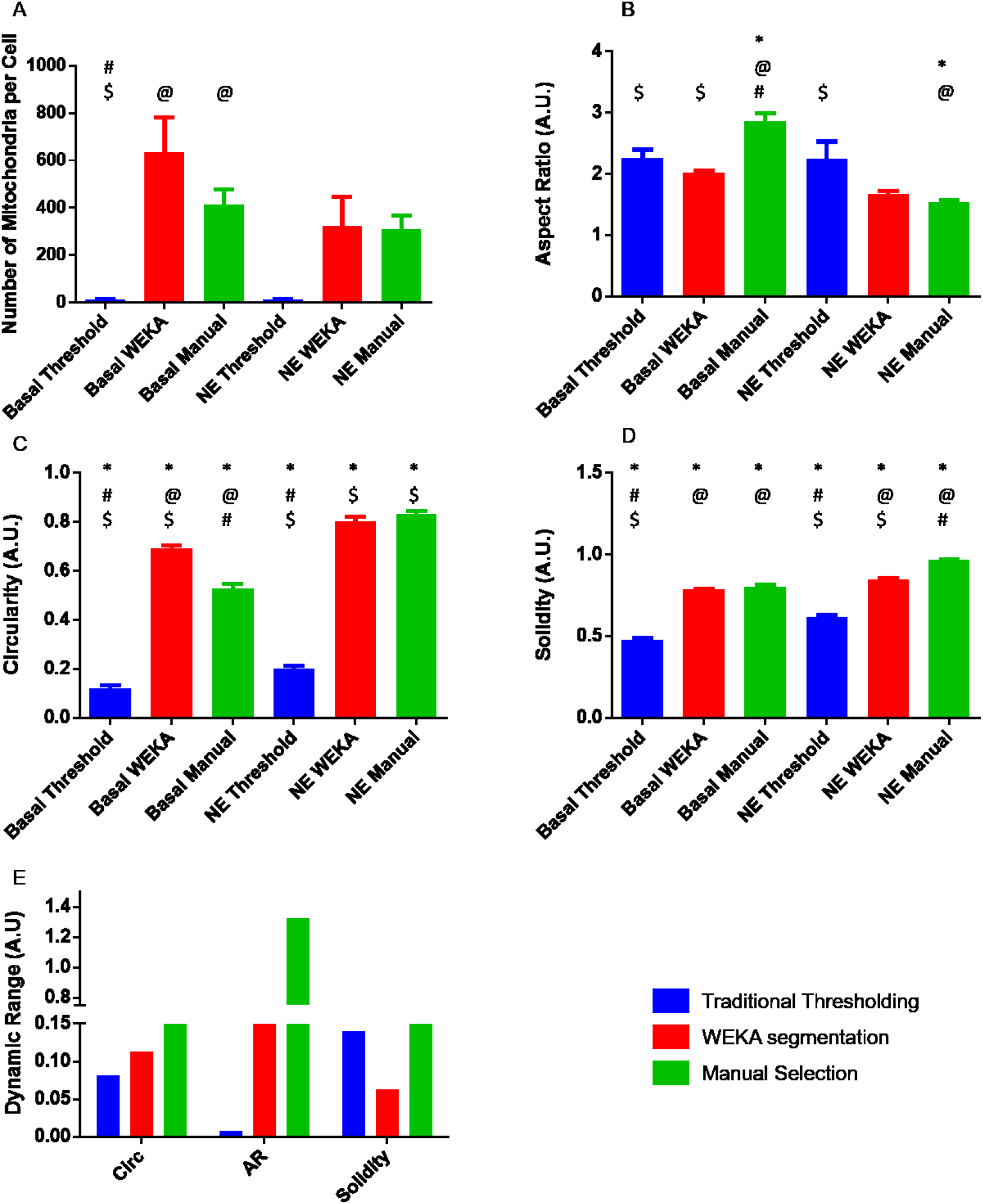
WEKA segmentation detects mitochondrial fragmentation in NE-activated BAT. A clear trend was exhibited by traditional thresholding to detect fewer mitochondrial objects per cell due to background fluorescence and airy haze surrounding mitochondrial objects in the images (A). WEKA segmentation detected significantly more mitochondrial objects than traditional thresholding but was not significantly different from manual ROI creation. The WEKA classifier more aggressively splits long filaments than a human analyst, leading to a trend of increased mitochondrial objects in the basal (nonstimulated) condition compared to manual analysis (A). This largely unavoidable bias is also evidenced by the significant decrease of AR with WEKA segmentation when compared to manual analysis (B). This artifact inherent to the algorithm necessitates a larger sample size for observed trends to be significant. However, Circ (C) and Solidity (D) exhibit significant increases with NE-activation of brown fat, suggesting reduced mitochondrial connectivity upon activation and validating the WEKA-based segmentation approach. WEKA segmentation increases dynamic range of measurements in both circularity and AR, error not shown because dynamic range was calculated from the averages shown in panels (A)-(D). (E) Each bar is the mean of >10 cells per condition from 3 separate experiments with error bars representing SEM; p≤0.05 by one-way ANOVA: (*) significant difference between basal vs NE conditions of same analysis method; (@) significant difference vs threshold analysis of same condition; (#) significant difference vs WEKA analysis of same condition; ($) significant difference vs manual analysis of same condition.

## DISCUSSION

Due to the increasing size (terabytes), quantity, and quality of data produced by digital microscopy in science, rapid, repeatable, and novel methods for processing these data are required. Given the difficulty of accurate image segmentation in service of unbiased measurements, we have created a novel segmentation method to observe subpopulations of mitochondria within cells. Segmentation of images in creative ways provides additional granular approaches in existing and newly and formerly acquired images, providing additional ways to extract as much data as possible from micrographs. This peridroplet analysis methodology may be expanded by others to address similar hypotheses: for instance, are lysosomes near the nucleus distinct from those at the cell periphery? By adapting the above approach to dilate the nuclear region instead of a lipid region, such an investigation is possible. Such investigations are rare (Leonard et al., 2015), partly because light microscopes only recently achieved sufficient resolution to facilitate this type of analysis (Weisshart, 2014). While the biological implications of this ROS formation we discovered remains to be elucidated, a recent publication suggests that cytosolic mitochondria are more oxidative, while the peridroplet mitochondria are essential for lipid droplet formation (Benador et al., 2018). The increased ROS formation observed here suggests that the enhanced oxidative capacity comes at the cost of potential oxidative damage. Additionally, a novel machine learning segmentation method for mitochondria more accurately segments mitochondria than traditional methods. By combining this segmentation protocol with a high-throughput approach to image analysis, we have designed a workflow that expedites an unbiased analysis of mitochondrial parameters. These tools will not only enable more time-and cost-effective research, but also provide a platform from which other scientists may improve and build better analysis protocols.

Subcellular objects pose a unique challenge in advanced light microscopy, partly due to the refraction limit of light microscopes (Abbe, 1873). Some tools, including the Airyscan detector from Zeiss (Weisshart, 2014), provide novel ways to approach super resolution light microscopy and further enhance the quality of micrographs. With these improvements in imaging technology, image analysis is becoming more approachable and more necessary. While the above WEKA method largely utilized super resolution micrographs, the recognition of subpopulations of mitochondria was performed on images acquired without super resolution techniques. In fact, the WEKA classifier is sufficiently accurate to segment mitochondria from non-super resolution micrographs, as demonstrated in Fig. 4. Indeed, the images analyzed in Fig. 6 were not super-resolution images; WEKA segmentation accurately detected mitochondrial fragmentation from these images.

The method contained herein has been validated on images from several different microscopes but primarily from a Zeiss LSM 880 Airyscan. All images were in focus and sufficiently magnified that a human could recognize individual mitochondria within the cytoplasm of the cell. This level of detail is generally acquired by an objective lens of 63X magnification or higher, with a NA of 1.4. All images were initially acquired with at least 1024 × 1024 pixels. The high resolution images permit detailed manual cropping of the cell from its field of view, a necessary step to ensure robust segmentation of mitochondria.

These tools are significantly more effective when the input micrographs are of high quality and resolution. Poor signal-to-noise ratio, low fluorescence in samples, and many other caveats of fluorescent microscopy influence the efficacy of image analysis. A recent publication details the ideal input image quality and common problems and artifacts encountered with analyses of this type (Harwig et al., 2018). For instance, out of focus images and images with resolutions less than 0.165 μm per px may not yield accurate results – the algorithms have not been validated at lower resolutions. Because the macro and protocol published here are designated for use with images collected by our lab, some tweaking may be required to use with images composed of different channel orders. Other common issues and their simple solutions have been published recently by another group (Harwig et al., 2018).

The protocols here provide tools to segment and quantify lipid droplet number, morphology, and intensity in brown adipocytes. We also describe a novel approach to segment mitochondria within these same cells in order to investigate mitochondria immediately adjacent to lipid droplets versus mitochondria in the rest of the cytosol. While such granulated approaches have been performed in the past (Glancy et al., 2015), they were undertaken in muscle tissue and revealed a novel energy conduction pathway along a mitochondrial network. Here we show a phenotypic difference between mitochondria immediately adjacent to the lipid droplet and the mitochondria distributed within the cytosol. Moreover, we provide a method for improved mitochondrial segmentation and measurement via the WEKA trainable segmentation plugin. Not only is this approach different from previous mitochondrial machine learning classification schemes (Leonard et al., 2015), but it provides an open and transparent tool with the potential to be refined and improved. The development of a WEKA classifier for mitochondrial segmentation expands the available input images significantly; a WEKA classifier is much better at segmenting poorly-acquired images than most conventional methods, preventing the issues mentioned above from excluding images from analysis. More importantly, input images from lower quality microscopes are generally accepted and properly segmented, assuming sufficient quality.

More detailed metadata of the images used in this method are contained in the methods. Brown adipocytes provide an ideal test case for these tools. Not only do the mitochondria in the cytosol differ from those surrounding lipid droplets (Benador et al., 2018), but also BAT mitochondria fragment upon norepinephrine-induced thermogenesis (Wikstrom et al., 2014), meaning accurate quantification of these phenomena can demonstrate the validity of our analysis protocols. This provides a new tool to analyze and accurately quantify mitochondrial parameters from images of insufficient resolution to apply conventional thresholding techniques, again as illustrated in Fig. 4. This increased flexibility and accuracy will provide more robust analyses within the field going forward.

## Supporting information

Supplemental Data - Macros and Classifier

