## Supplemental Data - Macros and Classifier for "High-Throughput Image Analysis of Lipid-Droplet-Bound Mitochondria": mito roGFP peridroplet and cytosolic mitos manual region check FINAL.ijm

```
//this macro asks user for threshold of cell body, then measures FI.
//it outputs the results to a .csv, and that can be opened in excel

//this is handy in EVERY macro
//it fixes the weird background changes that imagej does and also standardizes the overlays
run("Colors...", "foreground=white background=black selection=yellow");
run("Options...", "iterations=1 count=1 black edm=Overwrite");
//clear results table
run("Clear Results");

print ("Welcome to peridroplet/cytosolic mitochondrial analysis - BAT edition by Nathanael Miller."); 
print ("Please be sure you have a stacked image in the format Green-Red-BF-Purple");
dilations=getString("number of dilations to perform", 10);

rawdir = getDirectory("Choose the raw folder");
//savedir = getDirectory("choose the result folder");

//print("raw: " +rawdir);
//print("save: " +savedir);
list=getFileList(rawdir);

//set up batch processing

list=getFileList(rawdir);
for (i=0; i<list.length; i++)
{
	run("Bio-Formats Importer", "open=["+rawdir+list[i]+"] autoscale color_mode=Default view=Hyperstack stack_order=XYCZT");
	//this is the actual analysis procedure
	run("Split Channels");
	//close the BF image
	selectWindow("C3-"+list[i]);
	close();
	//store each image name by channel color for reference in this macro
	selectWindow("C2-"+list[i]);
	red=getTitle; 
	selectWindow("C1-"+list[i]);
	green=getTitle;
	selectWindow("C4-"+list[i]);
	purple=getTitle;
	//Begin peridroplet analysis
		//run("Threshold...");
		selectWindow(red);
		//begin nile red BG subtraction
			run("Duplicate...", " ");
			redBG=getTitle; 
			setAutoThreshold("Otsu dark");
			setOption("BlackBackground", true);
			run("Convert to Mask");
			run("Despeckle");
			run("Invert");
			run("Divide...", "value=255.000");
			//now you have a mask with 1 or 0 to multiply by original
			imageCalculator("Multiply create 32-bit", redBG, green);
			rename("greennodroplet");
			greennodroplet=getTitle;
			imageCalculator("Multiply create 32-bit", redBG, purple);
			rename("purplenodroplet");
			purplenodroplet=getTitle;
			selectWindow(greennodroplet);
			run("Duplicate...", " ");
			greencyto=getTitle;
			selectWindow(purplenodroplet);
			run("Duplicate...", " ");
			purplecyto=getTitle;
		//green and purple should have 0's in droplet regions
		selectWindow(red);
		setAutoThreshold("Otsu dark");
		setOption("BlackBackground", true);
		run("Convert to Mask");
		//this permits user defined number of dilations; each dilation is 1 pixel
		run("Despeckle");
		for(n=0; n<dilations; n++)
		{
			run("Dilate");
		}
		//run("Watershed");	
		//Sets the scale to 1 pixel = 1 pixel
		run("Set Scale...", "distance=1 known=1 pixel=1 unit=pixel global");
		//Defines the minimum area of ROIs.  Change the lower limit to define larger areas.
		run("Analyze Particles...", "size=50-Infinity pixel show=[Overlay Outlines] add in_situ");
		//this defines what measurements are output by the macro
		run("Set Measurements...", "area mean standard min perimeter integrated limit redirect=None decimal=3");
		//measure green image peridroplet mitos
		selectWindow(greennodroplet);
		waitForUser("confirm peridroplet recognition");
		//run("Threshold...");
		//setAutoThreshold("MinError dark");
		setThreshold(1, 100000);
		roiManager("Measure");
		//adds a column to the results table for image name, then inserts it the first row with results from that image
		//since this is a workaround, the rest of the regions from the same image show "0"
		//then a new image name marks where the regions from the new image are
		rois=roiManager("Count");
		setResult("image name", nResults-rois, "peridroplet-"+green);
		updateResults();
		//measure purple image peridroplet mitos	
		selectWindow(purplenodroplet);
		//run("Threshold...");
		//setAutoThreshold("MinError dark");
		setThreshold(1, 100000);
		roiManager("Measure");
		setResult("image name", nResults-rois, "peridroplet-"+purple);
		updateResults();
		//deletes all ROIs
		for(pooploop=0; pooploop<rois; pooploop++)
		{
			roiManager("Select", 0);
			roiManager("Delete");
		}
	//END PERIDROPLET ANALYSIS
	
	//Begin Cytosolic mito analysis
		//duplicate green and purple to make masks
		selectWindow(greencyto);
		run("Duplicate...", " ");
		greenmask=getTitle;
		//selectWindow(purple);
		//run("Duplicate...", " ");
		//purplemask=getTitle;
		//threshold green and purple images for masking
		selectWindow(greenmask);
		//run("Threshold..."); 
		setAutoThreshold("MinError dark");
		setOption("BlackBackground", true);
		run("Convert to Mask");
		//selectWindow(purplemask);
		//run("Threshold...");
		//setAutoThreshold("MinError dark");
		//setOption("BlackBackground", true);
		//run("Convert to Mask");
		//combine the green and purple images with AND operator to ensure both are present in the image
		//imageCalculator("AND create", greenmask, purplemask);
		//probesum=getTitle;
		//subtract peridroplet regions from cytosolic
		imageCalculator("Subtract create", greenmask,red);
		//run("Watershed");
		run("Analyze Particles...", "size=1000-Infinity show=[Overlay Outlines] add in_situ");
		//measure green image cytosolic mitos
		selectWindow(greencyto);
		//run("Threshold...");
		//setAutoThreshold("MinError dark");
		setThreshold(1, 100000);
		roiManager("Measure");
		//adds a column to the results table for image name, then inserts it the first row with results from that image
		//since this is a workaround, the rest of the regions from the same image show "0"
		//then a new image name marks where the regions from the new image are
		rois=roiManager("Count");
		setResult("image name", nResults-rois, "cyto-"+green);
		updateResults();
		//measure purple image cytosolic mitos	
		selectWindow(purplecyto);
		//run("Threshold...");
		//setAutoThreshold("MinError dark");
		setThreshold(1, 100000);
		roiManager("Measure");
		setResult("image name", nResults-rois, "cyto-"+purple);
		updateResults();
		//deletes all ROIs
		for(pooploop=0; pooploop<rois; pooploop++)
		{
			roiManager("Select", 0);
			roiManager("Delete");
		}
	//END CYTOSOLIC ANALYSIS
	run("Close All");	
}
waitForUser("Go to results table, click [File --> save as], then save where you want the data.  ***SAVE AS A .CSV FILE***, then click 'OK'");
```
