## Supplemental Data - Macros and Classifier for "High-Throughput Image Analysis of Lipid-Droplet-Bound Mitochondria": WEKA SEGMENTATION BASIC 1 channel manual threshold region check copypaste to exce.ijm

```
//this macro asks user for threshold of cell body, then measures FI.
//it outputs the results to a .csv, and that can be opened in excel

//this is handy in EVERY macro
//it fixes the weird background changes that imagej does and also standardizes the overlays
run("Colors...", "foreground=white background=black selection=yellow");
run("Options...", "iterations=1 count=1 black edm=Overwrite");
//clear results table
run("Clear Results");

print ("Welcome to mitochondrial analysis - by Nathanael Miller. WEKA segmentation protocol by Dane Wolf."); 
print ("This macro will segment your images using the trainable WEKA segmentation tool of FIJI");
print ("If you receive a Java exception error when running the WEKA section, you must do the following:");
print ("Help -> Update -> Manage Update Sites -> Check 'ImageScience'. Click 'Apply Changes', then restart FIJI.");
print ("You will have the opportunity to check the segmentation and delete any extraneous regions via the ROI manager.");
print ("If you cannot see the mitochondria when it asks you to check the regions, go to Image -> Adjust -> Window and Level -> Auto");  


rawdir = getDirectory("Choose the raw folder");
//savedir = getDirectory("choose the result folder");

//load classifier path as variable
classifier = File.openDialog("Open the classifier you want to use for WEKA segmentation")
//replace(classifier, \, "\\"); 
print (classifier);

//print("raw: " +rawdir);
//print("save: " +savedir);
list=getFileList(rawdir);

//set up batch processing
run("Set Measurements...", "area mean standard min perimeter shape integrated limit redirect=None decimal=3");

list=getFileList(rawdir);
for (i=0; i<list.length; i++)
{
	run("Bio-Formats Importer", "open=["+rawdir+list[i]+"] autoscale color_mode=Default view=Hyperstack stack_order=XYCZT");
	//this is the actual analysis procedure
		roiManager("reset")
		//make this "originaltiff" and then redefine the notiff string as "original" if imagej can't find the image name. the next 3 lines are designed to remove ".tif"
		originaltiff=getTitle;
		point=lengthOf(list[i])-4;
		original = substring(list[i], 0, point);
		run("Duplicate...", "duplicate channels=0");
		mitoseg=getTitle;
		//run("Subtract Background...", "rolling=50");
		//setTool("freeline");
		//call trainable weka segmentation plugin
		run("Trainable Weka Segmentation");
		wait(3000);
		selectWindow(mitoseg);
		selectWindow("Trainable Weka Segmentation v3.1.2");
		wait(3000);
		//load classifier into WEKA
		call("trainableSegmentation.Weka_Segmentation.loadClassifier", classifier);
		wait(3000);
		//THESE ARE THE OPERATIONS THAT YOU WANT WEKA TO DO! ONLY GETTING A PROBABILITY MAP
		selectWindow("Trainable Weka Segmentation v3.1.2");
		call("trainableSegmentation.Weka_Segmentation.getProbability");
		wait(3000);
		beep();
		//waitForUser("click OK when probability maps are generated");
		selectWindow("Probability maps");
		run("Stack to Images");
		selectWindow("Background");
		close();
		selectWindow("Mitochondria");
		//setThreshold(0.09, 1);
		//setAutoThreshold("Default dark");
		run("Threshold...");
		waitForUser("Set your desired threshold, then click OK");
		setOption("BlackBackground", false);
		run("Convert to Mask");
		run("Despeckle");
		run("Analyze Particles...", "size=0.1-Infinity show=Overlay add");
		selectWindow(originaltiff);
		roiManager("Show All");
		waitForUser("Check region recognition, then click OK");
		//run("Stack to Images");
		//remove ".tif" from original variable
		selectWindow(originaltiff);
		//rereference as "green" and measure
		green=getTitle;
		roiManager("Measure");
		rois=roiManager("Count");
		//updates results table with name of image just measured
		setResult("image name", nResults-rois, "green "+green);
		updateResults();
		//selectWindow("c:2/2 - "+original);
		//rereference as "red" and measure
		//red=getTitle;
		//roiManager("Measure");
		//updates results table with name of image just measured
		//rois=roiManager("Count");
		//setResult("image name", nResults-rois, "red "+red);
		//updateResults();
		beep();
		waitForUser("Copy the results table to a separate worksheet for each cell (ctrl-a, ctrl-c, ctrl-v), then click OK");
		run("Clear Results");
		
	run("Close All");	
}
//waitForUser("Go to results table, click [File --> save as], then save where you want the data.  ***SAVE AS A .CSV FILE***, then click 'OK'");
```
